# The evolution of social dominance through reinforcement learning

**DOI:** 10.1101/2020.07.23.218040

**Authors:** Olof Leimar

## Abstract

Groups of social animals are often organised into dominance hierarchies that are formed through pairwise interactions. There is much experimental data on hierarchies, examining such things as winner, loser, and bystander effects, as well as the linearity and replicability of hierarchies, but there is a lack evolutionary analyses of these basic observations. Here I present a game-theory model of hierarchy formation in which individuals adjust their aggressive behaviour towards other group members through reinforcement learning. Individual traits such as the tendency to generalise learning between interactions with different individuals, the rate of learning, and the initial tendency to be aggressive are genetically determined and can be tuned by evolution. I find that evolution favours individuals with high social competence, making use of individual recognition, bystander learning and, to a limited extent, generalising learned behaviour between opponents when adjusting their behaviour towards other group members. The results are in good agreement with experimental data, for instance in finding weaker winner effects compared to loser effects.

## Introduction

The empirical study of dominance hierarchies is nearly a century old (Schjelderup-Ebbe, 1922) and has identified important phenomena such as winner, loser, and bystander effects. Work in the field has also pointed to potential challenges for the understanding of hierarchy formation, for instance by noting that hierarchy replicability is sometimes low even though hierarchies are nearly linear (e.g. Chase et al., 2002).

A number of mathematical models of dominance hierarchy formation have been developed, both mechanistic and evolutionary models (reviewed by Mesterton-Gibbons et al., 2016). By assuming an updating of internal variables when individuals win or lose interactions (e.g. Dugatkin, 1997), or when they are bystanders to wins and losses by others (Dugatkin, 2001), mechanistic models have had some success in describing hierarchy formation, for instance in verifying that bystander effects promote hierarchy linearity. Because of the modelling complexities introduced by aggressive interactions between individuals that may differ in their fighting ability and previous experience, it has proven difficult to achieve evolutionary analyses of such mechanistic models. Nevertheless, winner and loser effects have been found in game-theory models by looking at simplified situations for groups of three individuals (Mesterton-Gibbons, 1999; van Doorn et al., 2003a,b).

To overcome these difficulties I here model the behavioural mechanisms of social hierarchy formation as reinforcement learning (e.g. Sutton and Barto, 2018). Recent work in neuroscience provides experimental support for the general idea that social dominance relations develop through processes that are similar to reinforcement learning (Kumaran et al., 2016; Ligneul et al., 2016; Qu et al., 2017; Zhou et al., 2018). Furthermore, observational learning, which is used as the mechanism underlying bystander effects in the model, is now part of the general neuroscience framework of reinforcement learning (Burke et al., 2010; Olsson et al., 2020). In addition to work in neuroscience and experimental psychology, there are thoroughly-studied reinforcement learning algorithms, with application in computer science and machine learning (Sutton and Barto, 2018), making the theory a good candidate for the integration of function and mechanism in game theory for biology (as discussed by McNamara and Leimar, 2020).

For evolutionary analysis of the model, the parameters of an individual’s learning rule are assumed to be genetically determined traits, and the evolution of these traits are studied through individual-based evolutionary simulations. An example of such a trait is an individual’s tendency to generalise its learned behaviour from one group member to other individuals, which gives rise to winner and loser effects during hierarchy formation. Other examples are learning rates, including bystander learning rates, and an individual’s tendency to be aggressive at the start of hierarchy formation. By performing such simulations for different sizes of social groups, as well as by keeping certain parameters at fixed values, which would correspond to evolutionary constraints, I investigate the evolution of social dominance behaviour in different contexts. This provides an understanding of the nature of social competence (Taborsky and Oliveira, 2012) for social dominance behaviour, by identifying the traits that are adaptive for an individual to adjust its behaviour during hierarchy formation. The analysis can reveal, for instance, whether individuals adapted to larger group sizes should show stronger winner and loser effects compared to those adapted to smaller group sizes, or under which circumstances winner effects should be weaker than loser effects.

The model is meant to be general, but still makes assumptions about cognitive capacities and how fitness depends on social interactions during a life history. As a possible example, one might consider interactions in groups of fowl (Schjelderup-Ebbe, 1922), in particular groups of male fowl (Pizzari and McDonald, 2019), in which individuals both display and fight during hierarchy formation.

To make contact with experimental work, I present model simulations illustrating potential results from experiments investigating winner, loser, and bystander effects, as well as experiments measuring hierarchy replicability and linearity. These results are reported both for evolved learning traits and for other values of some of these traits, such as the degree of generalisation and the bystander learning rate, in order to illustrate the influence of these traits in experiments. I also simulate an experimental set-up for estimating the amount of aggressive behaviour in interactions between a bystander and either the winner or loser of the bystander-observed interaction. The set-up tests the hypothesis that bystander effects differ between the cases, such that bystander effects are larger when a bystander interacts with the former winner (Earley and Dugatkin, 2002). I discuss whether the main model results are in qualitative agreement with observations of social dominance and I end by recapitulating the general game-theory approach used.

## Methods

### Model description

In each generation, there are a number of rounds or time steps of interaction, *t* = 1*, …, T*, and in each round there is a dominance interaction between two randomly selected individuals from a group of size *g_s_*. Each such interaction has the same general reward structure (see below), and group members can learn from the perceived rewards, which depend on their qualities *q_i_* (fighting abilities). At the start of a generation, individual qualities are randomly drawn from a normal distribution with mean *μ_q_* and standard deviation *σ*_1_, and they are assumed not to vary between rounds of the game. As a choice of zero point of the scale for *q*, let us assume that *μ_q_* = 0 (e.g., if fighting ability would be fully determined by size, *q* could be the logarithm of size, with the mean size in the population equal to one). Concerning what is ‘known’ by group members, assume that initially they do not have any particular information, including about their own fighting ability, but that they observe differences during dominance interactions and learn which actions to use through the rewards they perceive. The situation is then similar to traditional instrumental or operant conditioning.

For learning, the model uses the actor-critic method, described in section 13.5 of Sutton and Barto (2018). In this approach, an individual maintains and updates action preferences and estimates of values. The action preference (the ‘actor’) determines the behaviour to use in a given situation, which in the model is an interaction with a particular other individual, and the estimated value (the ‘critic’) is an estimate of the perceived reward from the interaction. The model makes a distinction between perceived rewards, which direct an individual’s learning, and evolutionary costs and benefits, which determine the action of natural selection. In addition to the description here, further details and explanation are found in Appendix A.

### Observations/states and actions

A real dominance interaction can consist of a sequence of several behaviours, including threat displays and fighting. The model simplifies this into two stages. In the first stage, interacting individuals, but not bystanders, make an ‘observation’ that acts like a state. Individuals observe some aspect *ξ* of relative fighting ability at the start of a dominance interaction, and also observe the opponent’s identity (individual recognition). The observation *ξ* might correspond to information from displays of some kind. The observation by each individual is statistically related to its quality (*q_i_*) and that of the opponent (*q_j_*). For the interaction between individuals *i* and *j* in round *t*, the observation is assumed to be

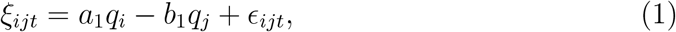

where *ϵ_ijt_* is an error of observation, assumed to be normal with mean zero and standard deviation *σ*. There is a similar expressions for *j* against *i*. By adjusting the parameters *σ*_1_ and *a*_1_, *b*_1_, and *σ* from equation (1) one can make the information about relative quality more or less accurate. The observation/state (*ξ_ij_, j*) is followed by a second stage, where individual *i* chooses an action, and similarly for individual *j*. The model simplifies to only two actions, A and S, corresponding to aggressive and submissive behaviour.

### Action preferences and estimated values

For an individual *i* interacting with *j* (with *j* ≠ *i*) in round *t*, *l_ijt_* denotes the preference for A. The probability that *i* uses A is then

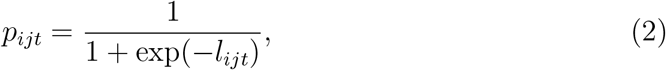

so that *l_ijt_* is the logit of the probability of using A. The model uses a linear (intercept and slope) representation of the effect of *ξ_ijt_* on the preference, and write *l_ijt_* as the sum of three components

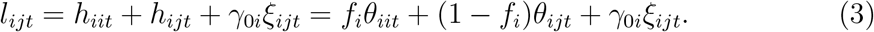

Here *h_iit_* = *f_i_θ_iit_* is a contribution from generalisation of learning from all dominance interactions, *h_ijt_* = (1 *− f_i_*)*θ_ijt_* is a contribution specifically from learning from interactions with a particular opponent *j*, and *γ*_0_*_i_ξ_ijt_* is a contribution from the current observation of relative quality. Note that for *f_i_* = 0 the learning about each opponent is a separate thing, with no generalisation between opponents, and for *f_i_* = 1 the intercept component of the action preference is the same for all opponents, so that effectively there is no individual recognition (although the observations *ξ_ijt_* could still differ between opponents). One can similarly write the estimated value 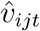 of an interaction as a sum of three components:

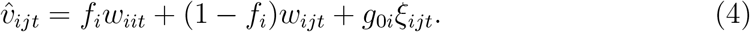

The actor-critic method updates *θ_iit_*, *θ_ijt_*, *w_iit_*, and *w_iit_* in these expressions based on perceived rewards, whereas *f_i_*, *γ*_0*i*_, and *g*_0_*_i_* are genetically determined.

### Exploration in learning

The trade-off between exploration and exploitation is a basic challenge for reinforcement learning algorithms (Sutton and Barto, 2018). For learning to be efficient over longer time spans there must be exploration (variation in actions), in order to discover beneficial actions. Learning algorithms, including the actor-critic method, might not provide sufficient exploration, because learning tends to respond to short-term rewards. In the model, exploration is implemented as follows: if the probability in equation (2) is less than 0.01 or greater than 0.99, the actual choice probabilities are assumed to stay within these limits, i.e. is 0.01 or 0.99, respectively. In principle the degree of exploration could be genetically determined and evolve to an equilibrium value, but for simplicity this is not implemented in the model.

### Perceived rewards

An SS interaction is assumed to have zero rewards, *R_ijt_* = *R_jit_* = 0. For an AS interaction, the aggressive individual *i* ‘wins’, which is associated with a perceived (primary) reward *R_ijt_* = *v_i_* for individual *i*, which is genetically determined and can evolve. The reward for the submissive individual *j* is zero, *R_jit_* = 0, and vice versa for SA interactions. If both individuals use A, some form of costly dominance display or fight ensues, with perceived costs (negative rewards or penalties) that are influenced by the qualities of the two individuals. The perceived rewards of an AA interaction are assumed to be

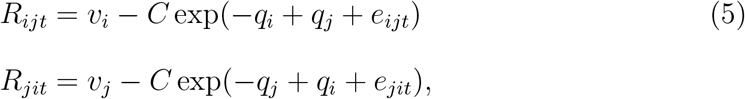

where *e_ijt_* is a normally distributed random influence on the perceived penalty, with mean zero and standard deviation *σ_p_*, and similarly for *e_jit_*. Note that it is not assumed that there is a ‘winner’ of the aggressive interaction. The reward components *v_i_* and *v_j_* are interpreted as innate motivation to perform aggressive behaviour. For simplicity, the parameter *C* is the same as for the evolutionary cost in equation (6) below. In effect, this gives a scale for the perceived rewards *v_i_* and *v_j_*.

### Evolutionary costs and benefits

The model assumes that there are evolutionary costs of AA interactions, because of injuries or exhaustion. For individual *i* interacting with *j* this is given by

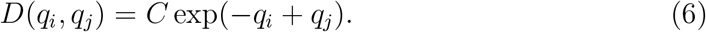

For simplicity, the dependence on the qualities has the same form as the perceived reward in equation (5). There are also evolutionary benefits from resources that become available (acquiring these resources is the ultimate function of dominance interactions). For simplicity, assume that resources are available after each interaction, and that the expected benefit for an individual that acquires the resource is *V*. If both individuals use A, the model assumes that they are preoccupied with aggressive behaviour, and neither gets the resource (it might for instance be that some other, unspecified agent gets the resource, or, if it is a potential mate, that the resource moves on). If one of a pair uses A and the other S, the aggressive individual is the winner and obtains the resource, and if both use S, they are equally likely to get the resource, or alternatively they share it (these give the same expected payoff).

### Learning updates

Let us first deal with updates for the individuals directly involved in a dominance interaction (see below for bystanders). The updates are driven by the prediction error (TD error)

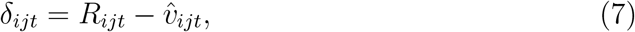

which is the difference between the actual perceived reward *R_ijt_* and the estimated value 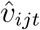. The learning updates for the *θ* parameters are given by

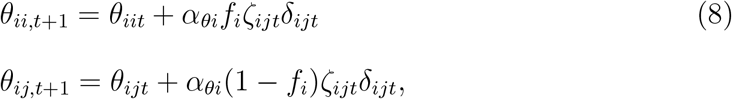

where

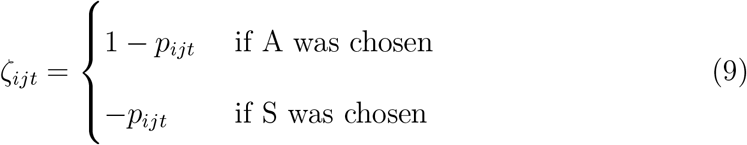

is referred to as a policy-gradient factor and *α_θi_* is the preference learning rate for individual *i*. Note that *ζ_ijt_* will be small if *p_ijt_* is close to one and individual *i* performed action A, which slows down learning, with a corresponding slowing down if *p_ijt_* is close to zero and S is chosen. There are also learning updates for the *w* parameters given by

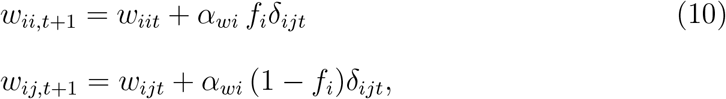

where *α_wi_* is the value learning rate for individual *i*. See SI for a derivation of these learning updates, using so-called policy gradient methods (following chapter 13 in Sutton and Barto, 2018). The values for the parameters at the start of a generation are also needed; let these be *θ_ii_*_0_ = *θ_ij_*_0_ = *θ*_0_*_i_* and *w_ii_*_0_ = *w_ij_*_0_ = *w*_0*i*_, where *θ*_0_*_i_* and *w*_0_*_i_* are assumed to be genetically determined. Note that *θ*_0_*_i_* determines how aggressive individual *i* is at the start of dominance hierarchy formation, which is an important aspect of social dominance behaviour.

### Bystander updates

To achieve a realistic description of social dominance, bystander effects are modelled as observational learning. In reinforcement learning theory, observation learning is conceptualised in a very similar way as other learning (Burke et al., 2010; Olsson et al., 2020), but without explicit specification of rewards. Instead, an observer can assign higher of lower significance to a particular event, implemented as the magnitude of an observational learning rate.

Applying this to dominance interactions, individuals other then the interacting pair *i* and *j*, for instance an individual *k*, could use the outcome to update the learning parameters. Assume that individual *k* only performs this updating if *i* and *j* use AS or SA (because there is no clear ‘winner’ in AA and SS interactions, and bystanders do not perceive the costs of AA interactions). The probabilities for individuals *i* and *j* to use A are *p_ijt_* and *p_jit_*, from equation (2). These are ‘true’ values and are not known by individual *k*. However, given that the outcome is either AS or SA, one readily derives that the logit of the probability that it is AS is

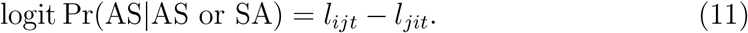

From equation (3) one can see that this involves various learning parameters for *i* and *j*. For bystander learning an assumption is needed about how an individual *k* represents this logit. A simple assumption is that *k* represents the logit as

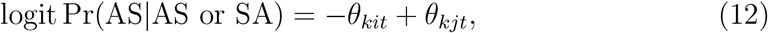

which entails that *k* does not use any information about *q_i_* or *q_j_*. The assumption is reasonable in that a large *θ_kit_* means that *k* behaves as if individual *i* is weak, and similarly for *θ_kit_*. Using the notation

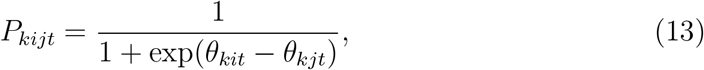

the bystander updates by *k* is assumed to be

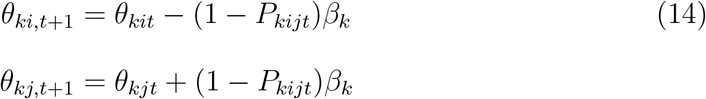

is *i* wins (outcome is AS) and

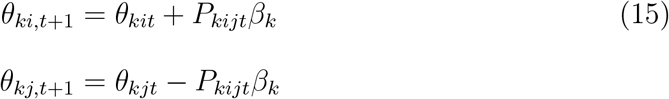

if *j* wins (outcome is SA). The parameter *β_k_* is a measure of how salient or significant a bystander observation is for individual *k*, and this parameter is assumed to be genetically determined and can evolve. These bystander updates are similar to the direct-learning updates of the actor component of the actor-critic model.

### Forgetting

Memory is an important cognitive capacity, which in practice will be more of less limited. Despite its importance, for simplicity the model does not implement a sophisticated memory mechanisms (including such things as interference between memories). Instead, we assume that there is a memory factor *m* such that the *θ* and *w* learning parameters decay towards their initial values with a rate *m* per time step.

## Evolutionary simulations

A number of parameters/traits in the model are assumed to be genetically determined and can evolve. The traits for an individual *i* are as follows: influence of states/observations on preference and value functions, from equations (3, 4), *f_i_*, *γ*_0*i*_, and *g*_0*i*_; perceived reward from performing A, from equation (5), *v_i_*; learning rates, from equations (8, 10, 14, 15), *α_θi_*, *α_wi_*, and *β_i_*; and initial values for the preference and value learning parameters, *θ*_0_*_i_* and *w*_0*i*_. In individual-based evolutionary simulations, each trait was implemented as determined by an unlinked diploid locus with additive alleles. For each generation, a population of *N* = 8000 individuals were randomly divided into groups of size *g_s_* equal to 4, 8, or 16. Each individual was assigned a quality *q_i_*, independently drawn from a normal distribution with mean zero and standard deviation *σ*_1_. Dominance-hierarchy formation then took place in each group, over a number of time steps such that each individual participated in an expected number of 200 dominance interactions. During a generation group members accumulated fitness payoffs as follows: a basic increment *V*_0_ per round of interaction, plus a payoff *V* for each resource acquired, and minus a cost as in equation (6) for each AA round. The next generation was formed by randomly selected parents from the entire population, with probabilities proportional to an individual’s accumulated payoff.

For each case reported in Table B1, simulations were performed over 5000 generations, repeated in sequence at least 100 times, to estimate mean and standard deviation of traits at an evolutionary equilibrium. These rather extensive simulations were performed to average out statistical fluctuations in the distribution of trait values over evolutionary time. As pointed out by, e.g., Bürger and Lande (1994), for mutation-drift-selection equilibrium dynamics under stabilising selection there can be substantial autocorrelation over generations of the population dynamical stochastic process.

### Standard parameter values

For several of the evolutionary simulations the following ‘standard values’ of parameters were used: fitness pay-offs, *V*_0_ = 0.5, *V* = 0.25, *C* = 0.2; distribution of individual quality variables, *a*_1_ = 0.707, *b*_1_ = 0.707, *σ*_1_ = 0.50; observations of relative quality, *σ* = 0.50; perceived penalty variation, *σ_p_* = 0.25; memory factor, *m* = 0.999. For these parameter values, around 50% of the variation in the observation *ξ_ij_* in equation (1) is due to variation in *q_i_ − q_j_*.

## Code availability

C++ source code for the individual-based simulations is available at GitHub, together with instructions for compilation on a Linux operating system: https://github.com/oleimar/socdom.

## Results

To illustrate the model (described in the Methods section), I present an example of the formation of a dominance hierarchy in a group of four individuals (Fig. 1). In the example, the learning traits are chosen to be at an approximate evolutionary equilibrium under the specified conditions, except that for simplicity there is no bystander learning in this example.

**Figure 1:**
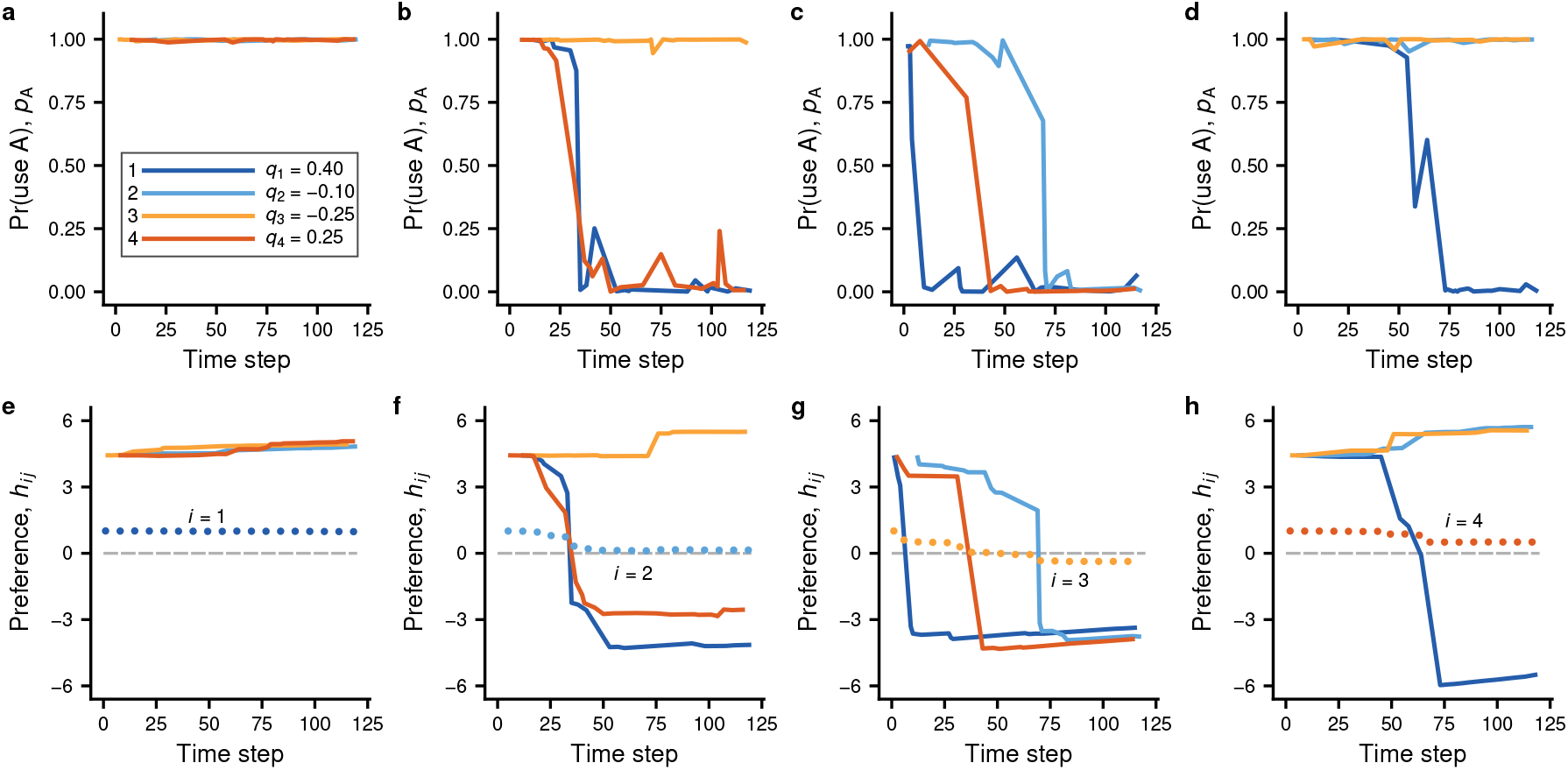
Illustration of the reinforcement-learning model for a group of size *g_s_* = 4, interacting over a total of *T* = 400 time steps. The first 120 time steps are shown, during which a stable dominance hierarchy is established. Panels (a) to (d) show the probability, from equation (2), of using the aggressive action A for individual *i* interacting with individual *j*, with one panel for each *i* and with colour coding for *j* as given in the inserted legend in panel (a). The inserted legend also gives the individual quality values *q_i_*. Panels (e) to (h) show the corresponding preference components *h_ij_*, from equation (3), with the colour-coded solid lines giving the component *h_ij_* for a specific opponent *j*, and the dotted lines showing the generalised component *h_ii_*, with one panel for each individual *i*. The parameter values for learning are as follows: *f_i_* = 0.19, *β_i_* = 0.0, *α_θi_* = 175.9, *α_wi_* = 0.09, *θ*_0*i*_ = 5.45, *w*_0_*_i_* = 0.002, *γ*_*0i*_ = 2.57, *g*_0_*_i_* = 0.025, and *v_i_* = 0.15. These are at an approximate equilibrium from evolutionary simulations, with *g_s_* = 4 and other parameters having the ‘standard values’ given in Methods (case 1 in Table B1 except that, for simplicity, bystander effects, from equation (15), are removed by putting *β_i_* = 0).

In each time step there is one round of dominance interaction between two randomly selected group members. In a round each individual first obtains an observation (with error) of the difference in fighting ability between itself and the opponent (not shown in Fig. 1; see equation (1)), for instance from display behaviours, and also observes the identity of the opponent. This is followed by a choice of actions, A (aggressive) or S (submissive), after which the round ends. An individual *i* perceives an internal reward *v_i_* from performing A, which can be thought of as a motivation to be aggressive. If both individuals choose A there is a fighting interaction, with perceived negative rewards (penalties) that depend on the fighting abilities (qualities) *q_i_* and *q_j_* of individuals *i* and *j* (equation (6)). From these each group member learns whether to use A or S against another group member. Panels (a) to (d) in Fig. 1 correspond to group members 1 to 4 and show the probabilities of using A against another group member (colour coded). Individual 1 is strongest and continues to almost always use A against the other group members throughout the time in the group (Fig. 1a). Individual 4 is the next strongest and initially tends to use A against all other group members, but eventually becomes submissive towards individual 1 (Fig. 1d). The other two individuals also start out aggressive but end up in lower dominance positions (Fig. 1b, c). The outcome is a linear hierarchy with dominance positions arranged according to fighting ability.

The model uses the actor-critic learning algorithm (Sutton and Barto, 2018). In this approach, the logit of an individual’s probability of using A is referred to as its preference for A and is expressed as a sum of components (equations (2, 3)). One could loosely compare with how effects are combined in logistic regression, but for the actor-critic method the preference components are changed through learning updates, driven by so-called prediction errors (equations (7, 8)). Two of the components are shown in panels (e) to (h) in Fig. 1; these correspond to panels (a) to (d). The solid lines (colour coded) show components specific to a particular opponent, whereas the dotted lines show the generalised component, which contributes to the preference for any opponent. Note that the opponent-specific components vary much more compared to the generalised component, and tend to change in an abrupt manner, producing rapid shifts from mostly aggressive to mostly submissive behaviour. A further illustration of the components over the entire time in the group appears in Fig. B1.

The reason for the large jumps in preferences seen in Fig. 1 is that the evolved preference learning rate is very high. Such high learning rates are a characteristic feature of the evolution of social dominance in the model. There are two learning rates in the model, a preference learning rate *α_θ_* and a value learning rate *α_w_*. The former is a parameter of the actor part of actor-critic learning and the latter belongs to the critic part; the critic learning rate does not evolve to high values. Comparing with real animals, the high preference learning rate could correspond to special cognitive processing of social dominance behaviour. Its function might be to allow rapid shifts from aggressive to submissive behaviour.

The degree of generalisation between opponents and the bystander learning rate are also important in social dominance, which is illustrated in Fig. 2. The learning traits are approximately at an evolutionary equilibrium (established by individual-based evolutionary simulations) for the cases in the figure. The comparison between blue and grey points show that suboptimally high generalisation (for the modelled situation) and a removal of bystander effects slows down learning about relative fighting ability and leads to a less sharp decrease in fights over time in the group. These effects are more pronounced in larger groups and they illustrate the nature of social competence in dominance-hierarchy formation, such as limited generalisation between opponents and substantial bystander learning rates, which are higher for individuals adapted to larger group sizes (see values of *β* for cases 1, 2, and 3 in Table B1). Pronounced generalisation between opponents, implying strong winner and loser effects, is detrimental to the formation of dominance hierarchies, as is illustrated in Fig. B2.

**Figure 2:**
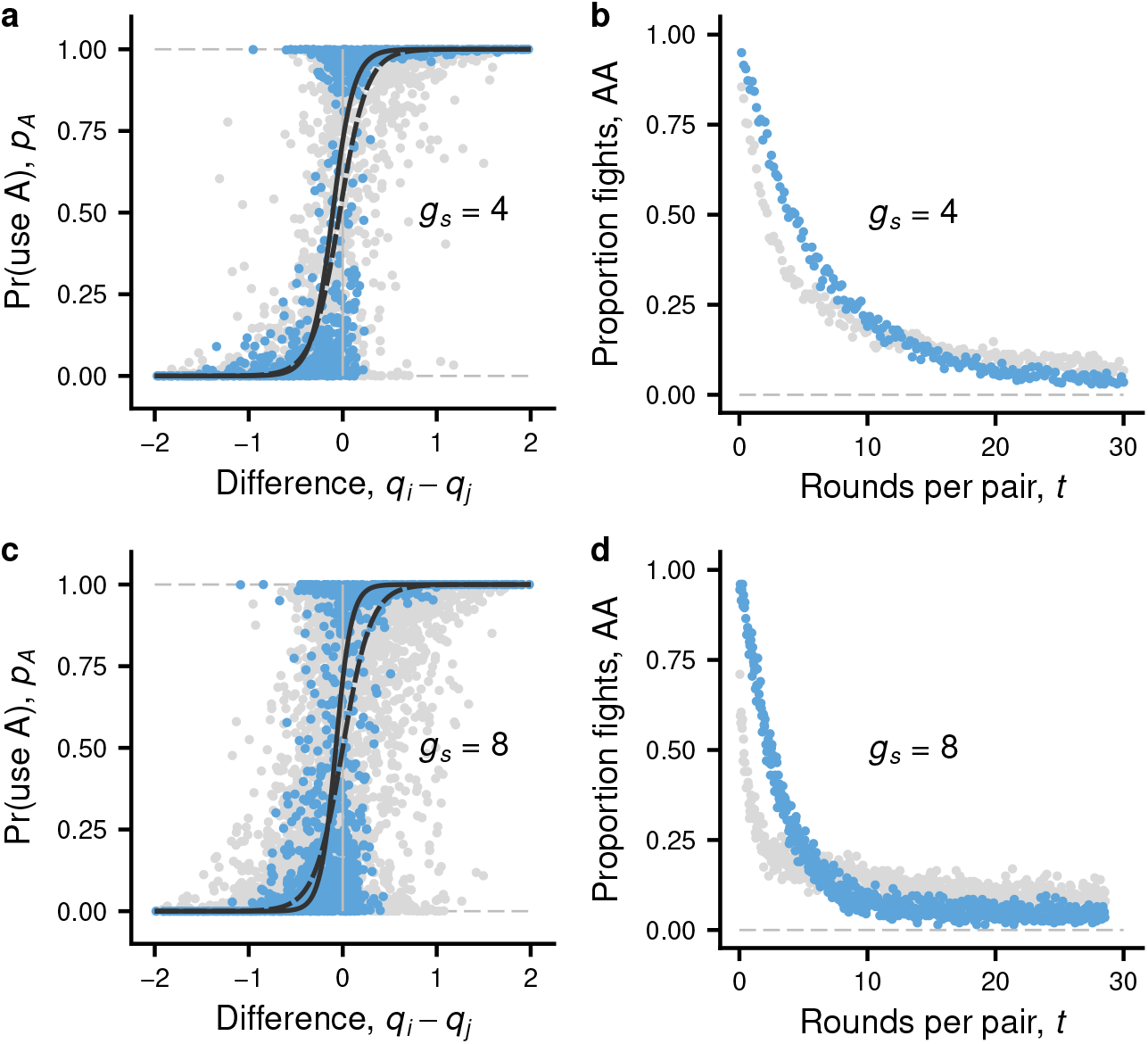
Influence of social competence on the learning about relative fighting ability and the decrease over time of aggression during hierarchy formation. Blue dots illustrate cases where the learning parameters, including the generalisation parameter (*f_i_*) and the bystander learning rate (*β_i_*), are at an approximate evolutionary equilibrium. The grey dots show the same except that generalisation is fixed at a suboptimally high value (*f_i_* = 0.5) and there is no bystander learning (*β_i_* = 0), corresponding to lower social competence. In panels (a) and (b), group size is *g_s_* = 4 with interactions over a total of *T* = 400 time steps, and in (c) and (d), *g_s_* = 8 with interactions over a total of *T* = 800 time steps. Panels (a) and (c) show the probability *p_ijt_* (equation (2)) of using the aggressive action A by an individual *i* plotted against the quality difference *q_i_ − q_j_* between itself and its opponent *j*, for times *t* corresponding to around 20 round per pair: each point gives the mean of *p_ijt_* over time steps *t* corresponding to 20 *±* 1 rounds per pair. The solid (for blue dots) and dashed (for grey dots) lines represent linear regressions on a logit scale. Panels (b) and (d) show the proportion of interactions that are AA (i.e. are fights) as a function of the number of rounds per pair. The dots overlap and represent averages for one simulated group over an interval of time steps. The evolved learning parameter values are given in Table B1, cases 1 and 7 for panels (a) and (b), and cases 2 and 8 for panels (c) and (d). Other parameters have the ‘standard values’ given in Methods.

For help in interpreting the evolved learning traits, Fig. 3 shows simulated experiments examining winner, loser, and bystander effects. In Fig. 3a, winner and loser effects are shown in darker blue for learning traits that evolved in groups of four, in red for different imposed values of the generalisation factor *f* (see equation (3)), and in light blue when the initial preference for A is kept at zero, corresponding to 50% chance of using A at the start of learning. As expected, the strength of winner and loser effects increases with the tendency to generalise between opponents (*f* = 0 means that there is no generalisation and *f* = 1 means that all opponents are treated as the same). Because of the experimental design, where the focal individual is tested against a matched opponent, there are substantial winner and loser effects for the evolved learning traits, although as seen from Fig. 1 the generalised preference component has rather little influence on hierarchy formation. The loser effects deviate more from *p*_A_ = 0.5 than the winner effects in Fig. 3a, except for the case of intermediate initial aggressiveness (light blue points), where the difference is small. This reveals a reason for weaker winner effects in the model. If individuals initially have a high preference for A, the actor learning updates from prediction errors while using A will be smaller than those from using S (equations (8, 9)), which can be compared with well-established results in the study of operant conditioning.

**Figure 3:**
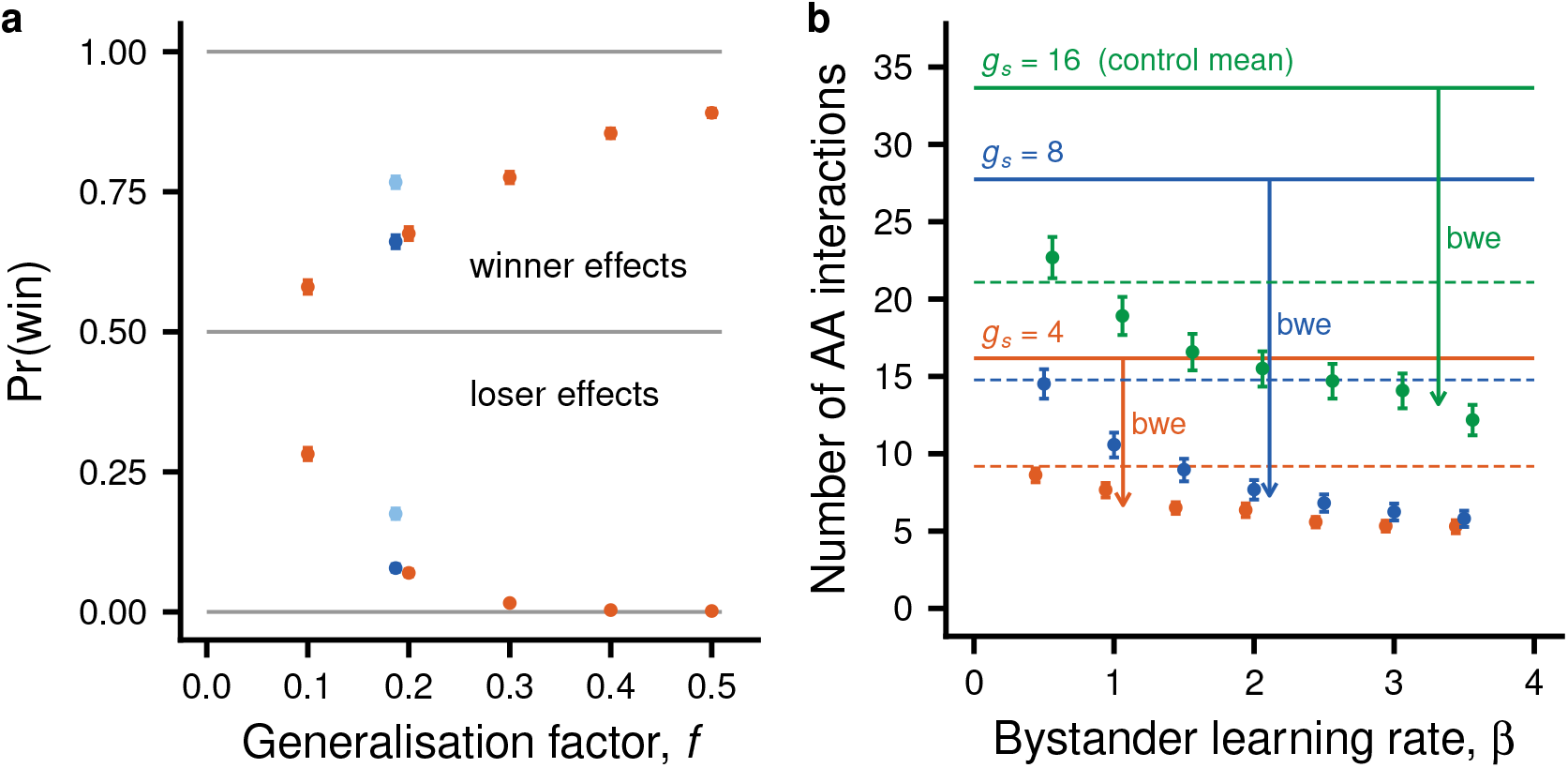
Winner, loser and bystander effects, as they might be observed in experiments. Panel (a) shows results from simulated winner- and loser-effect experiments. A focal individual either first experiences a win against a weaker or a loss against a stronger opponent (over at least 200 rounds and with Δ*q* = 0.5), and then interacts with another, matched opponent, scoring a win or a loss. The darker blue points (with 95% binomial confidence intervals) represent the probability of focal winning from 6400 replicated experiments of each kind, with learning parameters as in Fig. 2a (case 1 in Table B1). The red points illustrate the effects of changing the generalisation parameter *f_i_* from its evolutionary equilibrium, and the lighter blue points similarly illustrate setting the initial preference *θ*_0_*i* for action A to zero. Panel (b) shows results from simulated bystander-effect experiments, where a focal individual (with *q* = 0) first either observes or does not observe (control) a dominance interaction between a pair of individuals (over at least 10 rounds and with *q* = *−*0.25 and *q* = 0.25, respectively). The focal then interacts with the winner of the pair (solid lines and points), or the loser (dashed horizontal lines), and the number of rounds with fighting (AA rounds) is scored. The red, blue, and green horizontal lines (solid and dashed) show the overall mean for the control treatment, with different learning parameters (corresponding to cases 1, 2, and 3 in Table B1, with group sizes 4, 8 and 16, respectively). The vertical descending solid arrows show the bystander-winner effects (marked ‘bwe’) when the observer interacts with the winner, positioned at the evolved values of the by-stander learning rates *β_i_*. The red, blue, and green points (with 95% bootstrap confidence intervals) illustrate the consequences of changing the bystander learning rate, with 6400 replicates for each point. Other parameters have the ‘standard values’ given in Methods.

We can also compare with the extreme case of the grey points in Fig. B2c, d (case 11 in Table B1, with a negative *θ*_0*i*_), where there is no real hierarchy formation. For this case winner effects of the kind shown in Fig. 3 would be very strong (*p*_A_ = 0.99), as would be loser effects (*p*_A_ = 0.02). Such strong winner effects are not to be expected for individuals well adapted to life in social hierarchies.

The simulated experiments on bystander effects are inspired by previous experimental approaches (Johnsson and Åkerman, 1998; Earley and Dugatkin, 2002). The main result (Fig. 3b) is that bystander-winner effects, where a bystander meets the winner of an observed pair, are quite pronounced, and more so for individuals adapted to larger group sizes, while bystander-loser effects are small and not statistically detectable. Only the control results for bystander-loser effects are shown in Fig. 3b (dashed horizontal lines); the treatment results had confidence intervals overlapping the control, in spite of a very large sample size, so any bystander-loser effect in this example is very weak. Note also that the evolved bystander learning rates in Fig. 3b are higher when adapted to larger groups.

One way of investigating the influence of individual fighting abilities on dominance hierarchy formation is to let the same group form a hierarchy twice, with sufficient time in between so that individuals have no memory of the first at the second time (Chase et al., 2002). The simulated experiments of replicated hierarchy formation (Fig. 4a) show, as expected, that dominance relations are uncorrelated between replicates if there is very little variation in fighting ability *q*, but the correlation becomes high (but less than perfect) for the amount of variation that the learning traits are adapted to. There is little influence of bystander learning.

**Figure 4:**
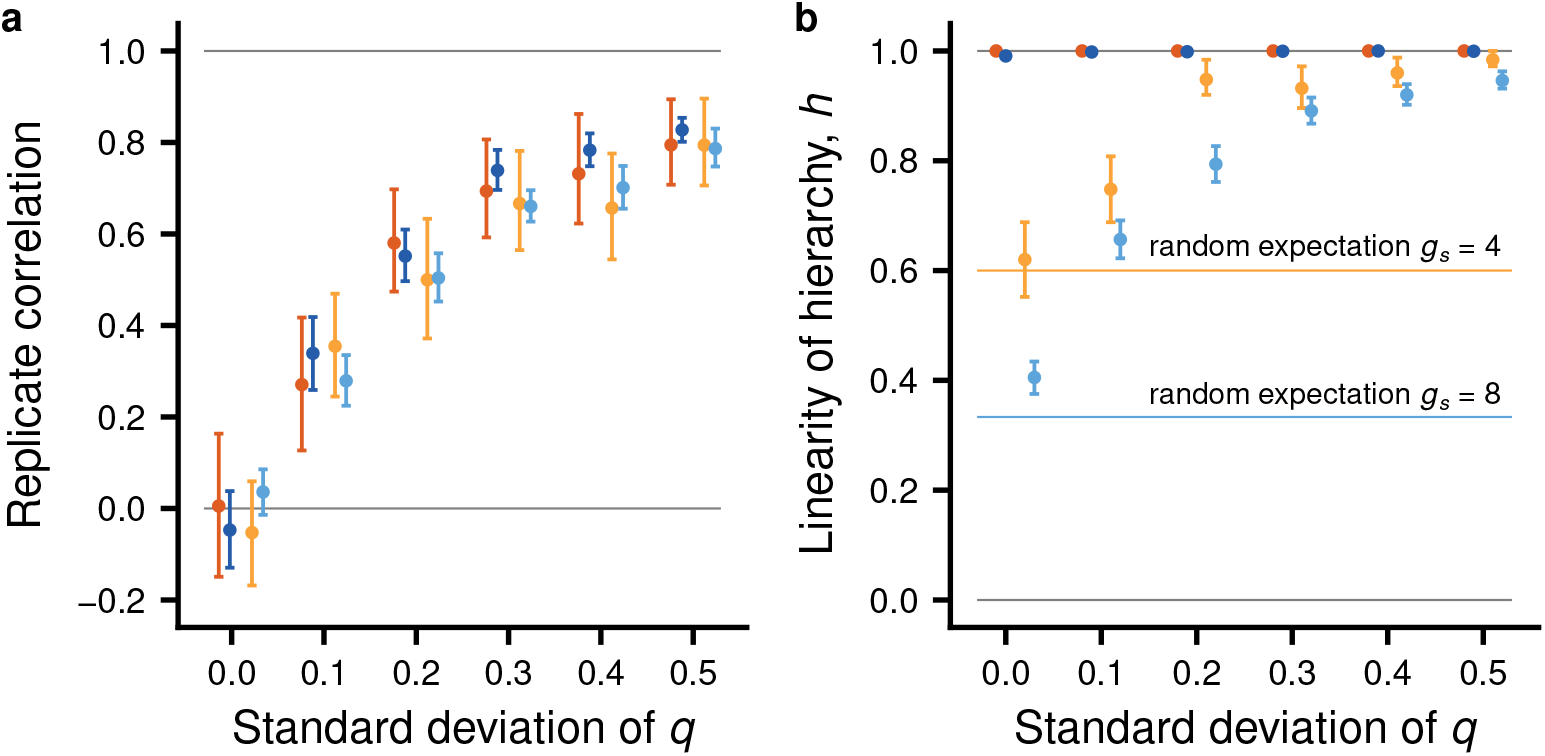
Replicability of dominance relations, in panel (a), and the linearity of dominance hierarchies, in panel (b), as a function the amount of variation in fighting ability in experimental populations, with and without bystander learning. The darker red and blue points (with 95% bootstrap confidence intervals) are for experiments with individuals with evolved learning parameters from cases 1 and 2 in Table B1, with group sizes *g_s_* = 4 and *g_s_* = 8, respectively. The orange and light blue points are for the corresponding evolved learning parameters when the bystander learning rate is fixed at *β_i_* = 0 (cases 4 and 5 in Table B1). In a simulated experiment, a group of size of either 4 or 8 is taken from an experimental population with some given SD of fighting ability *q* (values of SD: 0.01, 0.1, 0.2, 0.3, 0.4, 0.5; points are shifted left and right for clarity), ranging from nearly matched individuals in a group up to the variation in *q* to which the evolved learning parameters are adapted (i.e. an SD of 0.5). For each of these cases, and for each value of SD, the simulated data consists of 50 groups and 2 replicates per group. Group members are assumed to forget any learning between replicates. Panel (a) shows the correlation between replicates of the pairwise differences of the probabilities to use A from equation (2), *p_ij_ − p_ji_*, at the end of the time spent in the group (400 or 800 time steps). Panel (b) shows the Landau (1951) hierarchy index *h*, for which a value of 1 corresponds to a linear hierarchy. The orange and light blue horizontal lines indicate the expected values of the index for random dominance relations in groups of 4 and 8 individuals, respectively (given by eq. (29) in Landau, 1951). Other parameters have the ‘standard values’ given in Methods.

However, as seen in Fig. 4b, bystander learning has a major influence on hierarchy linearity Strikingly, when there is bystander learning, the linearity of hierarchies can be high even when replicates are essentially uncorrelated. Linearity is above random also without bystander learning (orange and light blue in Fig. 4b), but with the evolved bystander learning rates, linearity is essentially perfect both in groups of 4 and 8 individuals. To assess linearity the classical measure developed by Landau (1951) was used, taking into account the modifications by de Vries (1995) to account for possible ties.

In addition to the effects in Fig. 4, there are a number of other consequences of changing the amount of variation in fighting ability *q* in experimental populations. Many of these effects are quite intuitive, such as a stronger correlation between fighting ability and final dominance position when there is more variation in *q*, or more fighting interactions before hierarchy stability with less variation in *q*. For instance, for the cases in Fig. 4 of groups of size *g_s_* = 4 with bystander learning, the mean number of AA interactions per individual increases by a factor of 3, from around 21 to around 63, when the SD of *q* decreases from 0.5 to 0.01. For groups of size *g_s_* = 8 this effect is similar, with the mean number of AA interactions per individual increasing by a factor of 3, from around 29 to around 87, when the SD of *q* decreases from 0.5 to 0.01. It is thus more costly to settle dominance in groups with less variation in fighting ability.

## Discussion

By using the behavioural mechanisms of reinforcement learning, the model recovers and potentially explains several of the established aspects of dominance-hierarchy formation, such as winner, loser, and bystander effects. Among these are new explanations for why winner effects are weaker than loser effects and why bystander-winner effects are stronger than bystander-loser effects.

The parameter *f* in the model represents the strength of generalisation between opponents during hierarchy formation. This provides a link both to a long tradition in experimental psychology (Ghirlanda and Enquist, 2003) and to reinforcement learning theory (e.g. ch. 9 in Sutton and Barto, 2018). The parameter describes how a learning individual associates its experiences in dominance interactions with different characteristics of an opponent, like markings or colour patterns, ranging from those unique to each opponent (*f* = 0) to those shared by all opponents (*f* = 1). This aspect of the model thus explains winner and loser effects using the animal-psychology concept of stimulus generalisation. In principle this general idea could be tested experimentally through manipulation of characteristics mediating individual recognition in social groups (Tibbetts and Dale, 2007), examining if the availability of different characteristics influences winner and loser effects.

The model (Fig. 3a) provides a potentially general explanation of why winner effects are often found to be weaker than loser effects (Chase et al., 1994; Rutte et al., 2006; Oliveira et al., 2011). The explanation is that the magnitude of learning updates to the preference for A, given in equations (8, 9), depends on which action, A or S, is actually chosen. The update will be small if the action chosen had a high probability of being chosen. An interpretation is that the update is smaller when the choice is less ‘surprising’. This particular reason for a slowing down of learning appears not to have been tested experimentally, but it is in accordance with previous results and ideas about persistence in responding in operant conditioning, involving two-factor learning theory (Mowrer, 1951) and actor-critic learning theory (e.g. Maia, 2010). The explanation could be tested experimentally: the prediction would be that individuals showing high aggressiveness at the start of learning should show relatively weaker winner effects compared to loser effects.

The model implements observational learning (Burke et al., 2010; Olsson et al., 2020), sometimes referred to as vicarious reinforcement (Bandura, 1986), as a psychological mechanism underlying bystander effects. The bystander-learning updates, given in equations (14, 15), only change the opponent-specific components of the observer’s action preferences. These updates do not take into account information an observer might have about its own dominance status, which might be a simplification compared to what has been observed experimentally (Abril-de-Abreu et al., 2015).

The model simulations showed stronger bystander-winner than bystander-loser effects (Fig. 3b), which has been found in experiments (Earley and Dugatkin, 2002). The explanation for the model result is related to the explanation for stronger loser than winner effects. When a former observer interacts with the (stronger) winner of an observed pair, the former observer will have a somewhat reduced preference for A, and if S is used instead of A, the former observer rapidly learns to become submissive. On the other hand, when the former observer interacts with the (weaker) loser, the same explanation as the one above for a weak winner effect applies: just as in control conditions, the former observer has a strong tendency to be aggressive and does not change its preference for A to a noticeable degree.

The finding in Fig. 4, that hierarchies can be close to linear even when dominance relations are uncorrelated between replicates, is in accordance with experimental results (Chase et al., 2002). Note that, for the model, hierarchy linearity is a by-product of selection for traits influencing reinforcement learning; there is no direct selection for linearity. In the same way, for the model there is no direct selection for winner or loser effects – these are instead by-products of the evolved learning traits, including the strength of generalisation. Thus, the modelling does not assume that dominance-hierarchy formation can be decomposed into winner and loser effects of different strengths. It is even the case that winner and loser effects, which act through the generalised component of the action preference (Fig. 1e-d), can play a rather small role during hierarchy formation.

The simulated experiments illustrated in Fig. 4 showed that more fighting per capita, and thus greater cost, is required to establish dominance in groups with less variation in fighting ability. This is similar to the traditional idea/observation that it takes longer to settle contests with smaller differences in fighting ability. It is also more costly for individuals to settle dominance in larger groups. This prediction could easily be tested and is broadly consistent with findings on size-based hierarchies (e.g. Buston and Cant, 2006; Ang and Manica, 2010), that social groups that are large or contain individuals of similar size can become unstable because of costs of settling dominance relations.

The model is likely to be an oversimplification of any real case of dominance-hierarchy formation. The assumptions about evolutionary costs and benefits of dominance interactions, described in the Methods section, entail that reproductive success on average increases linearly with the dominance position, from lowest to highest. The effect of the dominance position on reproductive success might well vary between species and situations, and is often not empirically estimated in detail, but the model assumptions are at least broadly consistent with what is known about male fowl (McDonald et al., 2017; Pizzari and McDonald, 2019). The result for male fowl that experimentally induced wins or losses had rather little influence on subsequent hierarchy formation (Favati et al., 2017) is also broadly consistent with the model.

When comparing the model results with empirical observations, it is important to keep in mind that aggressive interactions, including those where winner and loser effects have been found, can represent adaptations to other situations than the formation of dominance hierarchies in social groups. Thus, contest over territories, or simply contests over resources that individuals encounter, often show effects of previous experiences of aggressive interactions (reviewed by Hsu et al., 2006). An example might be the strong winner effects found for a solitary parasitic wasp (Goubault and Decuignière, 2012) for which there are no social hierarchies, but where individuals can occur at high population densities and frequently contest resources. There are also model results dealing with effects of wins and losses in contests in large populations (e.g. Fawcett and Johnstone, 2010). Although there will be similarities between how the regulation of aggression becomes adapted to different situation, the current model applies specifically to the formation of hierarchies in social groups.

## Evolutionary modelling

By assuming several of the parameters of actor-critic learning to be genetically determined traits, one can study their evolution through individual-based simulation. In this way I found, for instance, that evolved rates of bystander learning are higher for individuals that are adapted to larger social group size (Table B1). The reason is that, in a lager group, each individual has proportionally fewer interactions with a particular other individual, making bystander learning more important. Compared to previous game-theory models of social dominance (Mesterton-Gibbons, 1999; van Doorn et al., 2003a,b), the current model is closer in spirit to previous mechanistic models (e.g. Bonabeau et al., 1996, 1999; Dugatkin, 1997, 2001).

Perhaps the most striking result from the evolutionary analysis is the extremely high evolved preference learning rates (Figs. 1, 2, Table B1). High rates of learning, including so-called one-trial learning, have for instance been found in taste aversion learning (Garcia et al., 1966; Shettleworth, 2010). For taste aversion learning the suggested explanation is that the perceived consequence (e.g. severe nausea) from ingesting unsuitable food is highly salient to the individual, and this is adaptive because individuals need to quickly learn to avoid poisonous or otherwise unsuitable food. There is not one-trial learning in social dominance, but the explanation for high rates of learning could be similar. For individuals to learn to form dominance relations in a reasonable amount of time, in particular to learn to be submissive for either of two closely matched individuals, large changes in the preference for A from fairly small prediction errors could be needed, which might explain the phenomenon. The high preference learning rates in the model could correspond to special cognitive (neural) processing of dominance interactions for real animals, which is in accordance with recent experimental results on social dominance (Kumaran et al., 2016; Ligneul et al., 2016; Qu et al., 2017; Zhou et al., 2018). This possible link between social dominance and reinforcement learning is one reason to think that the model could be relevant to social animals.

The general approach to modelling of social interactions used here is to introduce plausible behavioural mechanisms, and to treat parameters of these mechanisms as traits that can be tuned by evolution. As discussed by McNamara and Leimar (2020), this approach stands in contrast to much traditional game theory in biology, where an individual’s information about the environment, including about strategies of social partners, is represented as probability distributions over unknown aspects the environment. Traditionally in game theory, individuals use Bayesian updates, such as from prior to posterior distributions, to take into account information that is obtained. This can be referred to as a ‘small-worlds’ approach, because it is realistic mainly when an animal’s world, including the decision-making machinery of social partners, in some way is small and simple. The real world is a large world, especially for animals that interact and gain information about others over time. In such situations the traditional small-worlds approach becomes too challenging for modellers to achieve. More importantly, evolution is unlikely to have implemented an enormously complex decision-making machinery that properly performs the required Bayesian operations on probability distributions in high-dimensional spaces. Instead, evolution has lead to ‘large-worlds’ adaptations, in the form of behavioural mechanisms that work reasonably well in a range of circumstances. From this point of view, game-theory models of social behaviour ought to implement mechanisms that have strong empirical support, such as those from animal psychology and neuroscience (McNamara and Leimar, 2020).

Social hierarchies could be an area for which the large-worlds approach is particularly suited, by allowing stronger contact between modelling and observation. Using this approach here I have been able to outline important aspects of social competence for social dominance behaviour. These include individual recognition, good memory of previous interactions, some but rather limited generalisation between opponents, very high rates of preference learning, and efficient bystander learning. In the end it will be experiments that decide whether the mechanisms in the model are realistic, or if they must be changed to correspond to real dominance hierarchy formation.

## Acknowledgements

I thank John McNamara and Redouan Bshary for helpful comments. This work was supported by a grant (2018-03772) from the Swedish Research Council to the author.

## Appendix A Model details

This section contains further information and derivations, with the aim of explaining how the model relates to the broader treatment of reinforcement learning as presented by, e.g., Sutton and Barto (2018). As mentioned in the main text, the model considers the special case of only two actions, A and S. Another important simplification is that each round of interaction is assumed to be a so-called episode, meaning that there is no discounting of rewards over several time periods. Instead, the rewards (and penalties) from an interaction are determined by evolved characteristics, i.e. they are so-called primary rewards. This means that an individual does not learn that being dominant in an interaction is rewarding: the trait *v_i_* is an innate, evolved characteristic of the individual. What an individual learns in the model is instead whether to use A or S in interactions with a particular other individual.

Each round of interaction then starts with an observation or state, consisting of the observation *ξ_ij_* from equation (1) and the identity of the other individual. The model follows established knowledge about individual recognition (Tibbetts and Dale, 2007) in assuming that recognition is mediated by certain traits, for instance a pattern of colouration on the head or body of another individual. In order to introduce the crucial phenomenon of generalisation between stimuli, the model makes use of the general idea of function approximation for preference and value functions as developed in chapter 9 of Sutton and Barto (2018), in particular the linear methods. In this approach a state (i.e. another individual in the model) is represented using a number of features. The weights on the features of a state, such as the *θ* and *w* parameters in the model, express the required function approximation. Different states can share features to a greater or lesser degree, and a common approach is to use basis functions, such as Gaussian functions to express similarity between continuously varying stimulus components (referred to as radial basis functions by Sutton and Barto, 2018). This is similar to how generalisation is often described in animal psychology (Ghirlanda and Enquist, 2003). The model simplifies generalisation into two categories of traits: those that are shared by all opponents and those that are unique to a particular opponent. The generalisation factor *f_i_* in equations (3, 4) describes how strongly individual *i* takes into account the former in comparison to the latter when responding to opponents. The generalisation factor is assumed to be genetically determined, so the extent of generalisation can be tuned by evolution.

## Actor-critic learning updates

The principle used to formulate learning updates for the actor-critic method (see chapter 13 in Sutton and Barto, 2018) is to modify parameters *θ* and *w* of preference and value functions such that the expected reward from an episode is increased, which in the model corresponds to the expected reward from a round of dominance interaction. The actor-critic method implements this as approximate gradient ascent and is a policy-gradient method, inspired by the so-called policy-gradient theorem (see chapter 13 of Sutton and Barto, 2018). The updates to the policy parameters *θ* involves derivatives of the logarithm of the probability of choosing an action with respect to the parameters. Using equation (2), one can introduce the notation

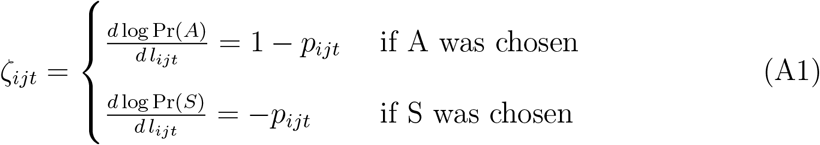

for the derivative of the logarithm of the probability of choosing an action, A or S, with respect to the preference for A, which corresponds to equation (9). The term policy-gradient factor can be used for *ζ_it_*, which is related to the concept of eligibility of the performed action for learning, as used in the study of actor-critic methods (Barto et al., 1983; Sutton and Barto, 2018). From equation (3) it follows that

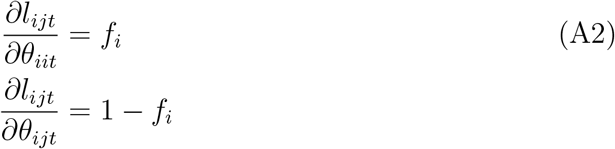

and from this one obtains the learning updates of the *θ* parameters as in equation (8). The actor-critic method also specifies updates of the *w* parameters of the value function. From equation (4) it follows that

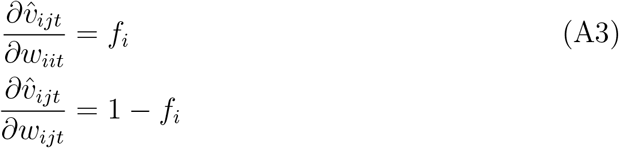

and from this one obtains the learning updates of the *w* parameters as in equation (10). These learning updates correspond to the one-step actor-critic updates described in chapter 13 of Sutton and Barto (2018), applied to the model.

## Appendix B Additional results

Figure B1 complements Fig. 1 in the main text by showing the dynamics of the preference components over the entire 400 time steps of the interaction. As seen in Fig. B1, there are occasional and rather large jumps in the components. They occur when an individual happens to select another action than the one indicated by its current action preferences.

**Figure B1:**
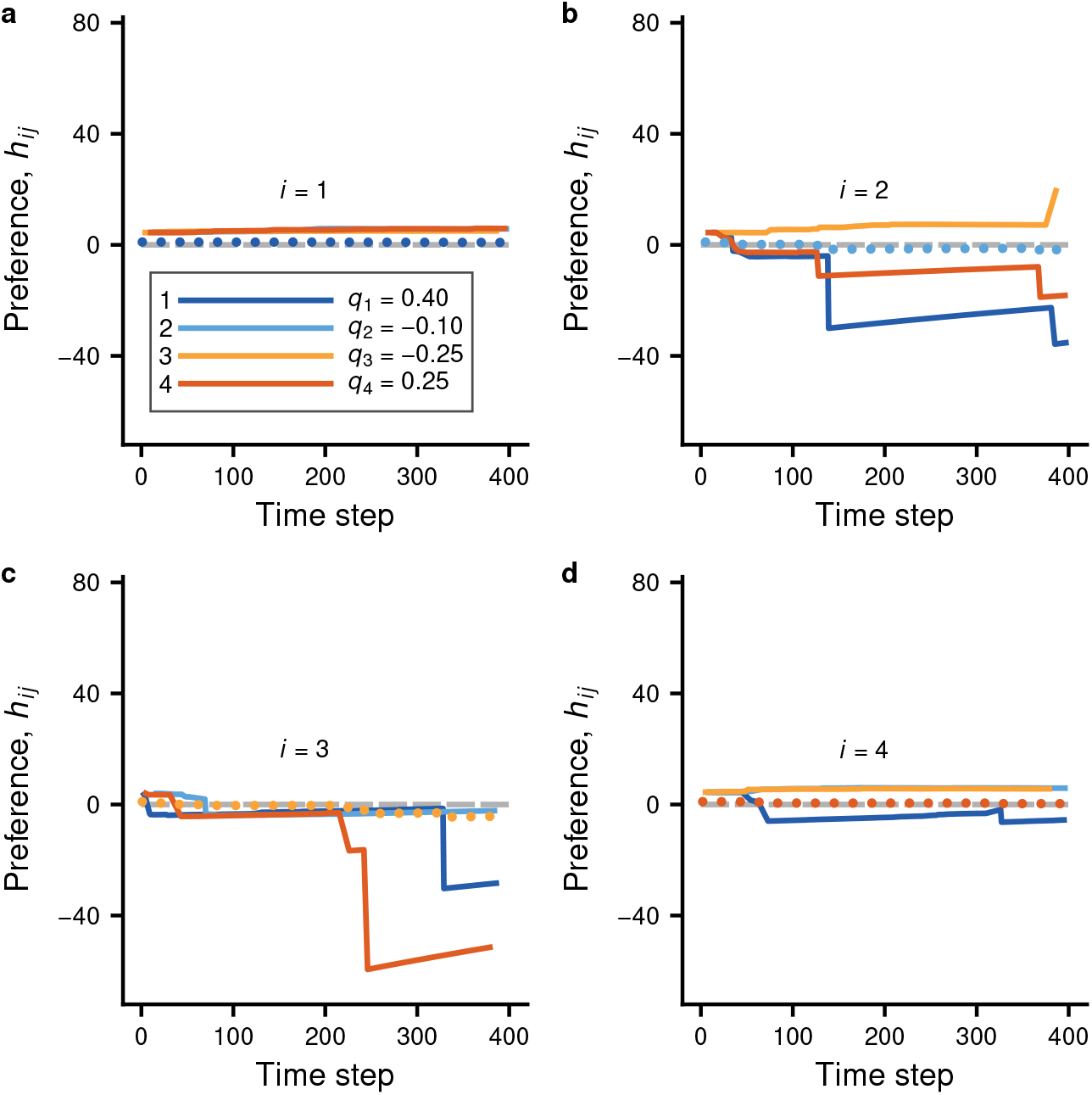
The panels (a) to (d) correspond to panels (e) to (h) in Fig. 1, but cover the entire 400 time steps of the dominance interactions and show a much greater range of preference values. The panels show the preference components *h_ij_*, from equation (3), with the colour-coded solid lines giving the component *h_ij_* for a specific opponent *j*, and the dotted lines showing the generalised component *h_ii_*, with one panel for each individual *i*. The inserted legend also gives the individual quality values *q_i_*. The big jumps in the preference components, as seen in the panels, occur when an individual ‘explores’ by choosing a different action (e.g. A instead of S). The parameter values for learning are as follows: *f_i_* = 0.19, *β_i_* = 0.0, *α_θi_* = 175.9, *α_wi_* = 0.09, *θ*_0_*_i_* = 5.45, *w*_0_*_i_* = 0.002, *γ*_0_*_i_* = 2.57, *g*_0_*_i_* = 0.025, and *v_i_* = 0.15.

Such deviations represent exploration by individuals, and they important for reinforcement learning. The consequence of exploration is typically that the current behavioural tendencies of an individual become stronger, in the sense that the jumps typically go in a direction that enhances an emerging dominance ranking.

Figure B2 shows a comparison like the one in Fig. 2 in the main text, but for two different cases, without bystander effects.

**Figure B2:**
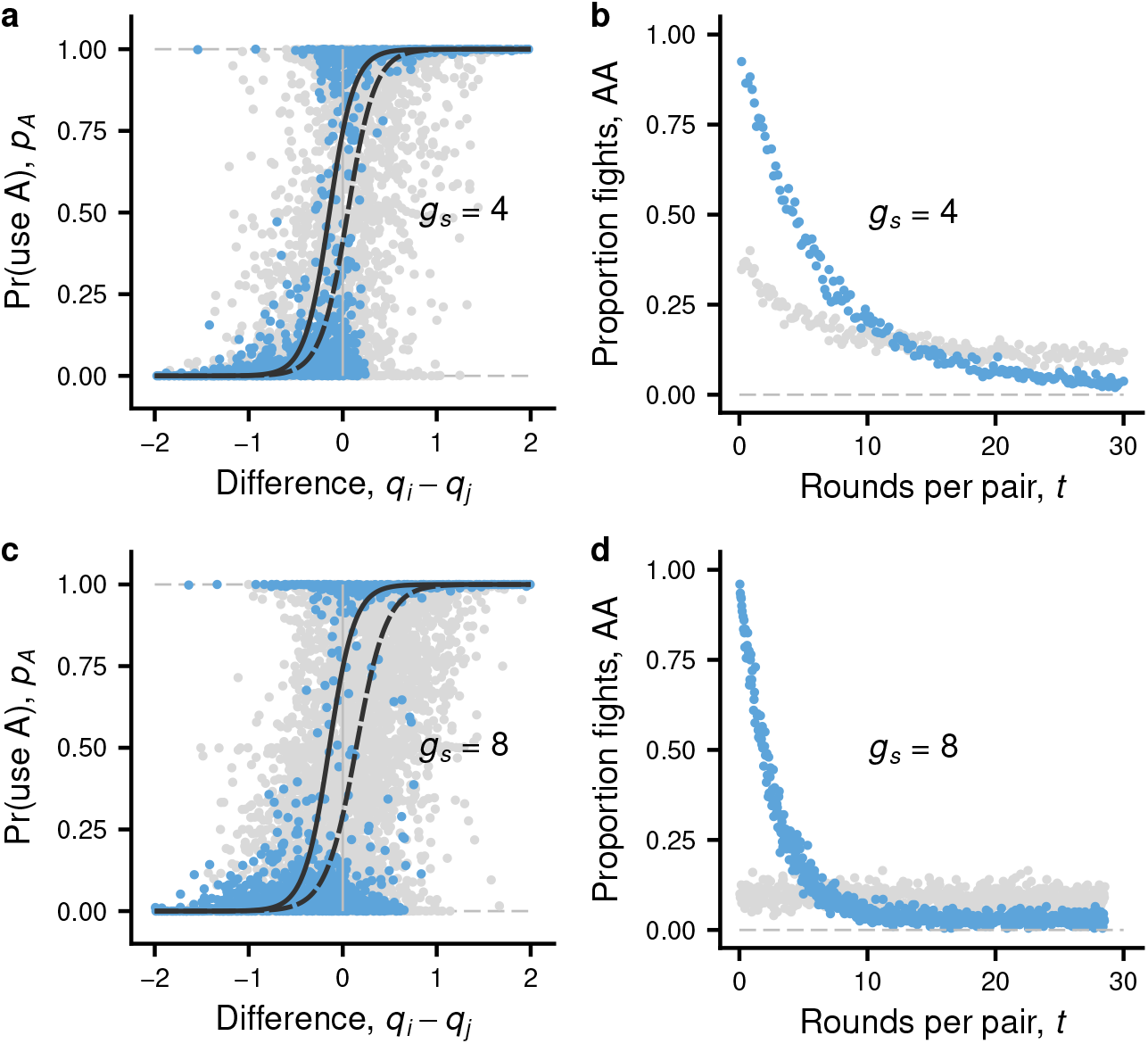
Influence of social competence on the learning about relative fighting ability and the decrease over time of aggression during hierarchy formation. There are no by-stander effects (*β_i_* = 0). Blue dots illustrate cases where the other learning parameters, including the generalisation parameter (*f_i_*), are at an approximate evolutionary equilibrium. The grey dots show the same except that generalisation is fixed at its maximal value (*f_i_* = 1), in effect preventing individual recognition, entailing very low social competence. Other details are as in Fig. 2. The learning parameter values are given in Table B1, cases 4 and 10 for panels (a) and (b), and cases 5 and 11 for panels (c) and (d).

For the grey dots in Fig. B2c, d (case 11 in Table B1), note that individuals do not learn to adjust their behaviour and thereby form a dominance hierarchy, but instead tend to use observations, as in equation (1), to settle interactions.

**Table B1:** Parameter values for 9 different cases of individual-based evolutionary simulations of social dominance interactions.

| case | $g_s$ | $T$ | $f_i$ | $\beta_i$ | $\alpha_{\theta i}$ | $\alpha_{wi}$ |
| --- | --- | --- | --- | --- | --- | --- |
| 1 | 4 | 400 | $0.19 \pm 0.01$ | $1.12 \pm 0.27$ | $175.9 \pm 20.0$ | $0.09 \pm 0.01$ |
| 2 | 8 | 800 | $0.16 \pm 0.01$ | $2.09 \pm 0.32$ | $428.2 \pm 37.4$ | $0.18 \pm 0.02$ |
| 3 | 16 | 1600 | $0.14 \pm 0.01$ | $3.26 \pm 0.20$ | $704.0 \pm 25.6$ | $0.31 \pm 0.04$ |
| 4 | 4 | 400 | $0.10 \pm 0.02$ | 0.0 | $110.6 \pm 11.3$ | $0.06 \pm 0.01$ |
| 5 | 8 | 800 | $0.07 \pm 0.01$ | 0.0 | $440.9 \pm 42.1$ | $0.11 \pm 0.01$ |
| 6 | 16 | 1600 | $0.06 \pm 0.01$ | 0.0 | $956.4 \pm 27.7$ | $0.21 \pm 0.04$ |
| 7 | 4 | 400 | 0.5 | 0.0 | $65.8 \pm 3.8$ | $0.21 \pm 0.04$ |
| 8 | 8 | 800 | 0.5 | 0.0 | $73.2 \pm 5.0$ | $0.37 \pm 0.05$ |
| 9 | 16 | 1600 | 0.5 | 0.0 | $74.3 \pm 9.2$ | $0.71 \pm 0.09$ |
| 10 | 4 | 400 | 1.0 | 0.0 | $23.8 \pm 2.5$ | $0.18 \pm 0.04$ |
| 11 | 8 | 800 | 1.0 | 0.0 | $21.8 \pm 3.4$ | $0.25 \pm 0.08$ |
| 12 | 16 | 1600 | 1.0 | 0.0 | $22.3 \pm 2.8$ | $0.34 \pm 0.12$ |

| case | $\theta_{0i}$ | $w_{0i}$ | $\gamma_{0i}$ | $g_{0i}$ | $v_i$ |
| --- | --- | --- | --- | --- | --- |
| 1 | $5.45 \pm 0.21$ | $0.002 \pm 0.004$ | $2.57 \pm 0.20$ | $0.03 \pm 0.004$ | $0.15 \pm 0.005$ |
| 2 | $5.98 \pm 0.17$ | $-0.01 \pm 0.005$ | $2.78 \pm 0.20$ | $0.03 \pm 0.004$ | $0.15 \pm 0.004$ |
| 3 | $5.67 \pm 0.24$ | $-0.04 \pm 0.007$ | $2.92 \pm 0.33$ | $0.03 \pm 0.005$ | $0.15 \pm 0.004$ |
| 4 | $4.99 \pm 0.21$ | $-0.01 \pm 0.004$ | $2.86 \pm 0.27$ | $0.03 \pm 0.004$ | $0.12 \pm 0.005$ |
| 5 | $5.05 \pm 0.15$ | $-0.03 \pm 0.004$ | $2.61 \pm 0.21$ | $0.02 \pm 0.003$ | $0.11 \pm 0.005$ |
| 6 | $4.52 \pm 0.27$ | $-0.06 \pm 0.005$ | $3.33 \pm 0.69$ | $0.02 \pm 0.003$ | $0.10 \pm 0.007$ |
| 7 | $3.60 \pm 0.12$ | $-0.01 \pm 0.009$ | $2.71 \pm 0.16$ | $0.03 \pm 0.004$ | $0.15 \pm 0.002$ |
| 8 | $2.63 \pm 0.18$ | $-0.06 \pm 0.016$ | $3.16 \pm 0.22$ | $0.04 \pm 0.005$ | $0.13 \pm 0.003$ |
| 9 | $-2.12 \pm 0.44$ | $-0.04 \pm 0.08$ | $12.59 \pm 1.10$ | $0.08 \pm 0.012$ | $0.10 \pm 0.007$ |
| 10 | $2.58 \pm 0.49$ | $-0.03 \pm 0.034$ | $8.58 \pm 0.66$ | $0.07 \pm 0.009$ | $0.09 \pm 0.004$ |
| 11 | $-2.87 \pm 0.56$ | $-0.02 \pm 0.075$ | $17.42 \pm 1.96$ | $0.09 \pm 0.015$ | $0.09 \pm 0.007$ |
| 12 | $-5.14 \pm 0.31$ | $0.05 \pm 0.123$ | $23.09 \pm 1.19$ | $0.11 \pm 0.020$ | $0.11 \pm 0.012$ |

## Notes

### Competing Interest Statement

The authors have declared no competing interest.

